# *Magnimaribacterales* marine bacteria genetically partition across the nearshore to open-ocean in the tropical Pacific Ocean

**DOI:** 10.1101/2025.06.17.660167

**Authors:** Oscar Ramfelt, Sarah J. Tucker, Kelle C. Freel, A. Murat Eren, Michael S. Rappé

## Abstract

The bacterial order *Magnimaribacterales*, previously known as the SAR86 lineage, is among the most abundant groups of planktonic bacteria inhabiting the global surface ocean. Despite their prevalence, our understanding of how this genetically diverse lineage partitions into units with coherent ecology and evolution remains limited. Here we surveyed multiple stations in the tropical Pacific Ocean using shotgun metagenomes and 16S rRNA gene amplicons to resolve distinct habitat preferences for *Magnimaribacterales* lineages across nearshore, offshore, and open-ocean environments. The comprehensive collection of genomes that captured a large fraction of the known evolutionary breadth of *Magnimaribacterales*, revealed patterns of ecotypic differentiation manifested primarily among genus-level clusters with specific clear preferences for distinct marine habitats. Enrichment analyses identified several functional genes associated with genomes from genera abundant in the nearshore environment, including those associated with sugar metabolism, peptide transport, and glycerophospholipid biosynthesis. Such metabolic adaptations likely facilitate the predominance of specific *Magnimaribacterales* genera in nearshore environments, promoting ecological partitioning across marine habitats.

**Importance:** Understanding the nature by which abundant, genetically-diverse planktonic marine bacteria organize into evolutionarily related and functionally coherent units remains an important question for scientists interested in the ecology of the global ocean. The bacterial order *Magnimaribacterales* (formerly SAR86) is one of the most prevalent clades in the global surface ocean, yet its ecological differentiation remains poorly resolved. By integrating metagenomic and amplicon analyses across a nearshore-to-open-ocean gradient in the tropical Pacific, this study reveals distinct habitat preferences within this lineage. The study also highlights how differences in distribution appear are primarily found between genus-level groupings. Our findings also highlight key functional traits, including sugar metabolism and glycerophospholipid biosynthesis, that were associated with the partitioning of nearshore and open-ocean *Magnimaribacterales*. This work enhances our understanding of the evolutionary processes shaping the diversity of one of the ocean’s most abundant bacterial clades.

## Introduction

The *Magnimaribacterales*, formerly known as the SAR86 clade (1), is an order-level lineage of marine bacteria within the *Gammaproteobacteria* that is ubiquitous within planktonic microbial communities of the global surface ocean (2, 3). Similar to the abundant *Pelagibacterales* (SAR11) lineage of *Alphaproteobacteria*, cells of the *Magnimaribacterales* have long been assumed to play an important role in the global carbon cycle because of their high cellular abundance in seawater (2, 4). However, despite sharing similar coarse features such as small cell size, small genomes and low GC content, analyses of gene repertoires for transporters and metabolic pathways reveal that *Magnimaribacterales* bacteria appear to avoid direct competition with co-occurring populations of *Pelagibacterales* by utilizing a distinct subset of the seawater organic carbon pool for heterotrophic growth. This includes a variety of fatty acid-containing molecules such as lipids, steroid-like polycyclic rings, and aromatic compounds (2, 3, 5). The *Pelagibacterales*, on the other hand, are known to generally degrade low-molecular weight compounds of the dissolved organic matter (DOM) pool that experience rapid turnover times (6).

In the case of the *Pelagibacterales*, integrative ’omics approaches have yielded a range of ecological and evolutionary insights. These include evidence of genetic differentiation into distinct clusters across ocean depths, along with genetic changes likely supporting this divergence (7–9). Additional distinct patterns of differentiation have also been observed across broad oceanic provinces, where comparative analyses reveal genetic variation among geographically separated populations (10, 11). Notably, differentiation is also apparent over finer spatial scales including between nearshore and open-ocean environments where putative functional differences between populations have been observed (12, 13). Temporal differentiation has also been reported, including seasonal shifts among closely related clades at the Bermuda Atlantic Time-series Study (BATS) site, suggesting a dynamic response to seasonality in the open-ocean (14, 15).

In contrast, studies of *Magnimaribacterales* ecology have primarily relied on whole-genome analyses and phylogenetic markers like the 16S rRNA gene (3, 5, 16–18). However, the limited phylogenetic resolution of the 16S rRNA gene, combined with sparse spatiotemporal coverage of available metagenomes, has left much of the ecology of *Magnimaribacterales* and the functional determinants of its niche partitioning explored. For instance, while some studies have suggested genetic differences between coastal and open-ocean *Magnimaribacterales* based on 16S rRNA sequences, these comparisons remain limited in scope (16, 19).

Here we take advantage of the Kāneʻohe Bay Time-series (KByT), a long-term study established in 2017 along the windward coast of the Hawaiian island of Oʻahu, to explore how *Magnimaribacterales* populations genetically and functionally differentiate across a stark nearshore to open-ocean environmental gradient (12, 20). KByT leverages easy access from nearshore to open-ocean waters in the tropical Pacific, serving as a natural laboratory to investigate the ecotypic differentiation of marine microorganisms (12). Sampled near-monthly, KByT spans a marine *Synechococcus*-dominated nearshore to *Prochlorococcus*-dominated offshore gradient (12, 20). In addition, the proximity of KByT to Station ALOHA, located ∼144 km to the north of Kāneʻohe Bay, allows us to directly link our observations with those of the Hawaiʻi Ocean Time-series (HOT) program within the North Pacific Subtropical Gyre (21, 22). Our findings are supported by leveraging extensive DNA sequence and environmental datasets from the surface ocean of both KByT and HOT, including 16S rRNA gene amplicons and shotgun metagenomes, providing a detailed view of how the *Magnimaribacterales* partitions across this gradient.

## Materials and Methods

### Sample description

Seawater samples were collected as part of KByT as previously described (12, 20). Briefly, starting in August 2017, approximately monthly samples were collected from a depth of two meters across ten stations spanning the nearshore environment within Kāneʻohe Bay on the island of Oʻahu, Hawaiʻi, to the offshore environment outside of the bay (Figure 1). In addition to samples filtered for analyses of microbial nucleic acids and phytoplankton pigments, a variety of other measures used to characterize the environment were obtained (Table S1). Thirty-six metagenomes were sequenced from samples collected between 2017-2021 (Table S1; 20, 23).

**Figure 1.**
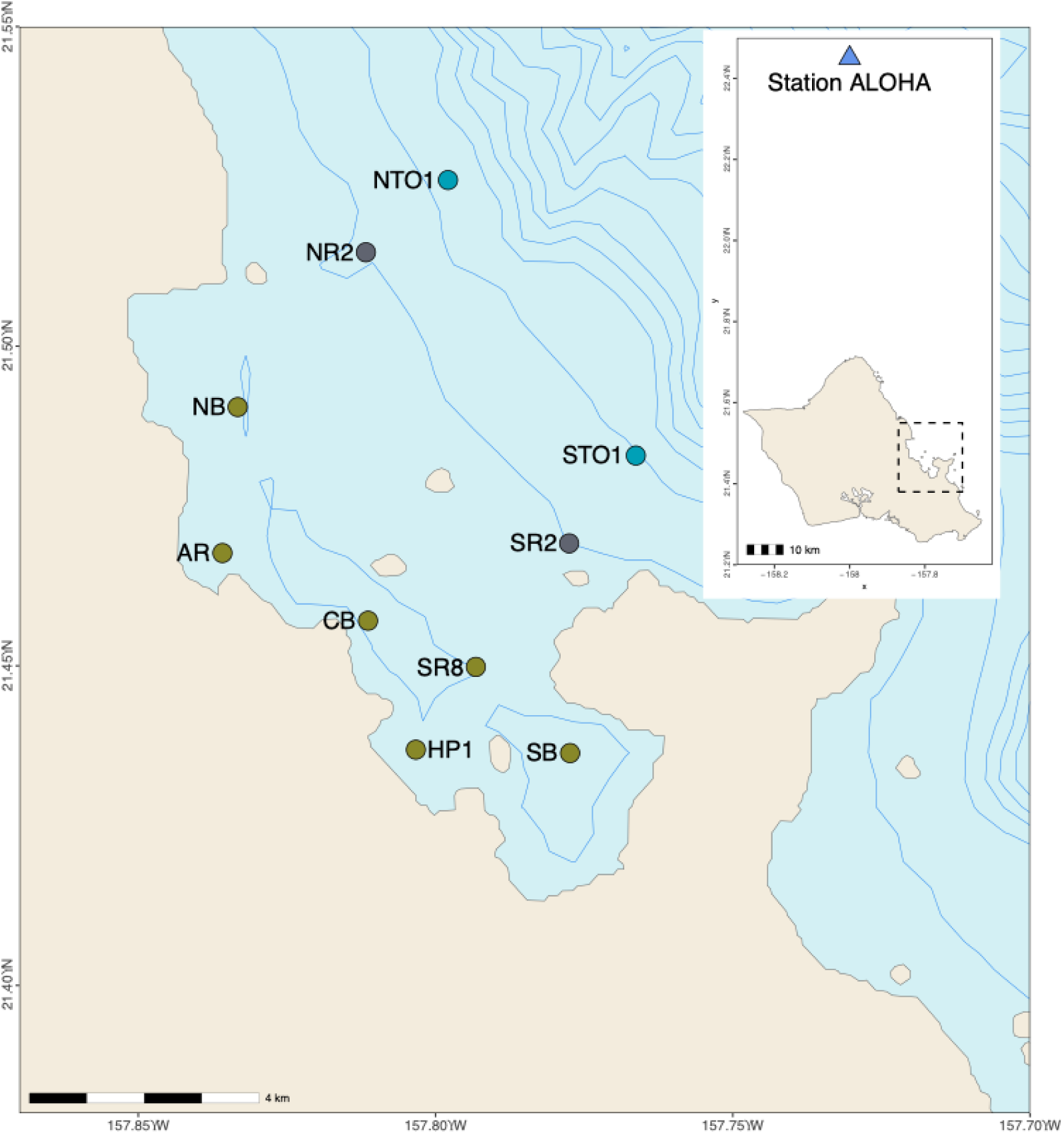
Map of Kāneʻohe Bay Time-series (KByT) sampling stations on the windward side of the Hawaiian island of Oʻahu. Background shading of station markers indicates environmental category (yellow - coastal; grey - transition; blue - offshore). The inset shows the location of Station ALOHA relative to Oʻahu.

Previously published metagenomes from surface ocean (0 - 75 m) samples collected as part of the Hawaii Ocean Time-series (HOT) at Station ALOHA in the North Pacific Subtropical Gyre (NPSG) were also included in these analyses (Table S1; 24). In addition to metagenome sequence data, we also re-analyzed a previously published 16S rRNA gene amplicon time-series survey from KByT samples, also from a depth of 2 m, spanning August 2017 to June 2019 (BioProject Accession PRJNA706753; 12).

### Ribosomal RNA gene amplicon analysis

Forward amplicon reads were imported into Qiime 2 v2022.11.1 (25), trimmed, and grouped into amplicon sequence variants (ASVs) using DADA2 v1.26.0 (26). ASVs found in only one sample or that had less than 10 reads across all samples in aggregate were excluded from further analyses. The taxonomy of each ASV was determined using the SILVA 99% OTUs full-length sequences dataset v138 (27) with the feature-classifier plugin (28). Features classified as part of the order “SAR86_clade” from the feature-classifier analysis were subsequently assumed to be a part of the *Magnimaribacterales* order.

A phylogenetic tree was constructed with the *Magnimaribacterales* ASV sequences and 16S rRNA genes extracted from the *Magnimaribacterales* reference genomes used in this study (n=64). Sina v1.7.2 (29) was used to align the *Magnimaribacterales* sequences with the Silva SSU Ref NR 99 v138.1 dataset and subsequently used to infer a phylogeny using RAxML v8.2.12 (30) with the GTRGAMMA model, 100 bootstraps, and a sequence alignment of 245 nucleotides. This tree was used to infer the family-level taxonomic classifications of each ASV (Figure S1). We replaced all ASV family classifications for f_SAR86_clade with the more recent family-level classifications from (3) using the “tools import” command in Qiime. These updated family classifications for *Magnimaribacterales* ASVs were used all downstream analyses with amplicons.

Relative abundance estimates for each ASV were calculated by first filtering out any sequences classified within the domain Eukaryote, domain Unassigned, and the Order Chloroplast. Then, any remaining ASVs had their relative abundance calculated using the ‘feature-table relative-frequency’ command from Qiime 2. To calculate family-level relative abundances the ‘taxa collapse’ command was used with a ‘–p-level’ of 5. This collapsed table was then used as the input to ‘feature-table relative-frequency’.

The filtered ASV table was used to estimate the cellular abundance of *Magnimaribacterales* by using the cellular abundances of non-chlorophyll containing cells presumed to be predominantly heterotrophic bacteria and archaea determined via flow cytometry from corresponding subsamples of the same seawater collection used for 16S rRNA gene amplicons (Table S1; 12). First, the dataset was further filtered by excluding any amplicon sequences related to the phylum Cyanobacteria from the previously filtered ASV table. The relative abundance of *Magnimaribacterales* within the reduced amplicon dataset was then multiplied by the cellular abundance of non-chlorophyll containing cells presumed to be predominantly heterotrophic bacteria and archaea (12).

### Metagenome assembly and binning

Five different co-assemblies were used to construct metagenome assembled genomes (MAGs) from shotgun sequenced bacterioplankton samples collected from KByT (Table S1). Three co-assemblies used metagenomes from samples collected between 2017-2019 and were grouped into nearshore, transition, and offshore categories based on previous analyses (KByT_1_Nearshore, KByT_1_Transition, and KByT_1_Offshore in Table S1; 12). Metagenomes collected between 2020-2021 were assembled separately and were split into nearshore and offshore groups based on the location of collection and co-assembled separately (KByT_2_Nearshore and KByT_2_Offshore in Table S1).

All metagenomes were quality filtered using ‘iu-filter-quality-minoche’ v2.12 (Table S2; 31). Quality filtered reads were assembled into contigs using MEGAHIT v1.2.9 (32), and contigs below 2500 base pairs in length were filtered out using ‘anvi-script-reformat-fasta’ v7.1 (33). Reads were mapped back to the filtered contigs using Bowtie2 (v2.3.5) (34) and were sorted and indexed using SAMTools v1.9 (35).

The filtered contigs and mapping results were used as input to create genome bins using the programs MaxBin v2.2.7 (36), MetaBAT v2.15 (37), and CONCOCT v1.1.0 (38). The program DAS Tool v1.1.4 (39) was then used to parse the results of the binning approaches, resulting in the bins used for downstream analyses.

Following binning, filtered contigs were used to create anvi’o contigs databases, and later converted to anvi’o profiles using ‘anvi-profile’. Coassemblies with multiple individual profiles were merged using ‘anvi-merge’. Bins identified from DAS Tool were imported into their respective profiles using ‘anvi-import-collection’. Dereplication of genomes was accomplished with dRep v3.3.0 with the -comp 50 option (39) and run twice; once with the 2017-2019 KByT genome bins as input and once with the 2020-2021 KByT genome bins as input. Genome bins were identified as *Magnimaribacterales* (order “SAR86” in GTDB-Tk) from the de-replicated genomes using the ‘classify_wf’ command of GTDB-Tk v1.7.0 (40) and were manually refined using the command ‘anvi-refine’ (41).

Post-refinement, the *Magnimaribacterales* genome bins were exported and quality checked using ‘taxonomy_wf’ within Checkm v1.2.0 (42) and a *Magnimaribacterales*-specific marker gene set (3). MAGs with a completeness <50% or contamination >7.5% were excluded from downstream analyses.

### Phylogenomics

A phylogenomic tree was constructed from an alignment of the *Magnimaribacterales* MAGs generated in this study with an exhaustive database of representative *Magnimaribacterales* genomes, including the genome of type strain *Magnimaribacter mokuoloeensis* str. HIMB1674 (3). The commands ‘identify’ and ‘align’ in GTDB-Tk v1.7.0 (40) were used to identify GTDB marker genes and create a multiple sequence alignment (MSA). Briefly, the ‘identify’ command used Prodigal v2.6.3 (43) to call genes and then HMMER v3.1b2 (44) identified the marker genes from the BAC120 gene set used by GTDB (45). The ‘align’ command used the output from ‘identify’ as input to create the MSA, which was then used as input by IQ-Tree v2.1.2 (46) for a maximum likelihood phylogenomic analysis using the LG + F + R7 model, which has been used previously for *Magnimaribacterales* phylogenomic analyses (3), and 1000 ultrafast bootstrap replicates (UFBoot) (47). A previously constructed taxonomic framework for the *Magnimaribacterales* that includes family and genus designations (3) was used as a scaffold from which to infer the taxonomy of new MAGs generated as part of this study. Visualizations of the phylogenetic tree alongside external data were constructed with ggtree v3.6.2 (48).

Average nucleotide identity (ANI) was calculated for all genome pairs using fastANI v1.32 (49), and subsequently used to group newly constructed MAGs into species clusters using previously described methods (50) and an ANI value of 93% previously used for the *Magnimaribacterales* (3). Representative genomes were selected from each species cluster using the following approach. If the species cluster contained a genome classified in (3) then that genome was used. If the species cluster consisted only of new MAGs assembled as part of this study then the MAG with the highest completion metric was used. This approach established 81 representative genomes that were used for read recruitment (“Representative” column in Table S3).

### Metagenomic read recruitment

The program Bowtie2 was used to map sequence reads from quality filtered KByT and Station ALOHA metagenomes to the representative genome set. SAMtools was used to sort and index the output sam files. Mapping metrics were calculated by importing bam files into anvi’o profiles using ‘anvi-profile’ and merged into a single profile with ‘anvi-merge’. Once merged, metrics for the merged profiles were exported using ‘anvi-summarize’.

The detection metric from anvi’o, which represents the proportion of a genome covered by reads, was used to help identify the presence of *Magnimaribacterales* genomes within metagenomes. Based on previous studies that have found detection values to follow a bimodal distribution (51), a detection value of 0.25 or greater was selected in line with other studies to identify the presence of a genome and avoiding false-positive signals (52). For any metagenomes where a genome had a detection level below the cutoff, its relative abundance in that metagenome was modified to zero. This was done to accurately reflect the absence of genomes from metagenomes where less than 25% of the genome recruited reads. This gave a matrix of presence/absence for each representative genome in each metagenome used in this study.

Using the matrix of the presence of each representative genome in each metagenome, it we then sought to identify the prevalence of each genome within the wider environments of nearshore, offshore, and open-ocean (Station ALOHA). The presence of a genome in an environment was measured as the proportion of metagenomes from each environmental grouping that the genome was found to be present in. As an example, if a genome was present in 4 out of 10 metagenomes from the nearshore environment it would be present in 40% of metagenomes.

To determine a suitable cutoff for whether a genome was present in an environment the ‘geom_density’ function within ggplot v3.4.2 (53) was used to make a smooth histogram for each environment. This showed that the offshore and open-ocean environment followed a bimodal distribution pattern with most genomes present in almost all metagenomes (near a 100%) or almost none (a value near 0%) in these two environments (Figure S2). In contrast, the nearshore environment, while displaying two peaks at 0% and 100%, showed two more subtle peaks at just over 75% and at around 60% (Figure S2). Based on these peaks the value of 50% was selected for a genome to be present in any environment meaning that if a genome was found in more than half of all metagenomes from an environment it would be considered present in that environment. Metagenomes originating from the transition environment of KByT were excluded from this analysis.

### Pangenomic and enrichment analysis

A pangenome was constructed from the *Magnimaribacterales* genome dataset using the anvi’o pangenomic pipeline (54). Anvi’o v8 (33) contig databases were generated for each genome using ‘anvi-gen-contigs-database’, which identified genes using Prodigal v2.6.3 (43). Genes were then annotated using clusters of orthologous genes (COGs) (55) and Kyoto’s Encyclopedia of Genes and Genome (56) via the commands ‘anvi-run-ncbi-cogs’ and ‘anvi-run-kegg-kofams’, respectively. Furthermore, EGGNOG annotations (database: v5.0.2) (57) were also determined by using the identified genes as input for the ‘emapper.py’ program v2.1.6 (58). These annotations were then imported to their respective contig databases using the ‘anvi-script-run-eggnog-mapper’ function of anvi’o. A genome-storage database was subsequently created using ‘anvi-gen-genome-storage’ and then the pangenomic analysis was run using ‘anvi-pan-genome’ with a mcl inflation value of 10 and a minbit value of 0.5. The ‘anvi-pan-genome’ command used DIAMOND v2.1.8 (59) to compare amino acid sequences between all genes found within the input genomes, and the Basic Local ALignment Search Tool (BLAST) v2.14.1 (60) from National Center for Biotechnology Information (NCBI) to identify gene clusters based on minbit values. The results of the pangenomic analysis were exported into a tabular format using ‘anvi-summarize’ for manual inspection.

To help identify functional enrichment among closely related genomes, the ‘anvi-compute-functional-enrichment-across-genomes’ pipeline was used (61). The pipeline was run twice for the *Magnimaribacterales* order using genus classifications as the grouping variable. The first run used KOfam annotations while the second run used annotations from COG. The results of the enrichment analysis were used to help identify differential gene distribution across environments.

## Results

### Suzuki family dominates the relative abundance of *Magnimaribacterales* 16S rRNA gene amplicons across KByT

Ribosomal RNA gene amplicon data showed that *Magnimaribacterales* constituted 2.0% to 4.3% of all amplicons, excluding those associated with Eukaryotes, chloroplasts, or unassigned sequences, across the three environments sampled through KByT (Figure 2A, Table S4). The Suzuki family was the most abundant of the four family-level lineages across all three environments, ranging from an average of 3.0% in the nearshore environment to 1.3% in the transition environment (Figure 2B, Table S4). Changes in cellular abundances were more pronounced, ranging from an average of 7.02 x10^4^ to 1.39 x10^4^ cells mL^-1^ in the nearshore and transition zones, respectively (Figure S3, Table S4). In addition to Suzuki, the CHAB-I-7 family also peaked in both average relative abundance and estimated absolute cellular abundance within the nearshore environment (Figure 2B, Figure S3, Table S4). While the RedeBAC7D11 and *Magnimaribacteraceae* families peaked in average relative abundance in the offshore environment (Figure 2B), the estimated absolute cellular abundance of *Magnimaribacteraceae* was actually higher in the nearshore environment compared to the offshore (Figure S3, Table S4).

**Figure 2.**
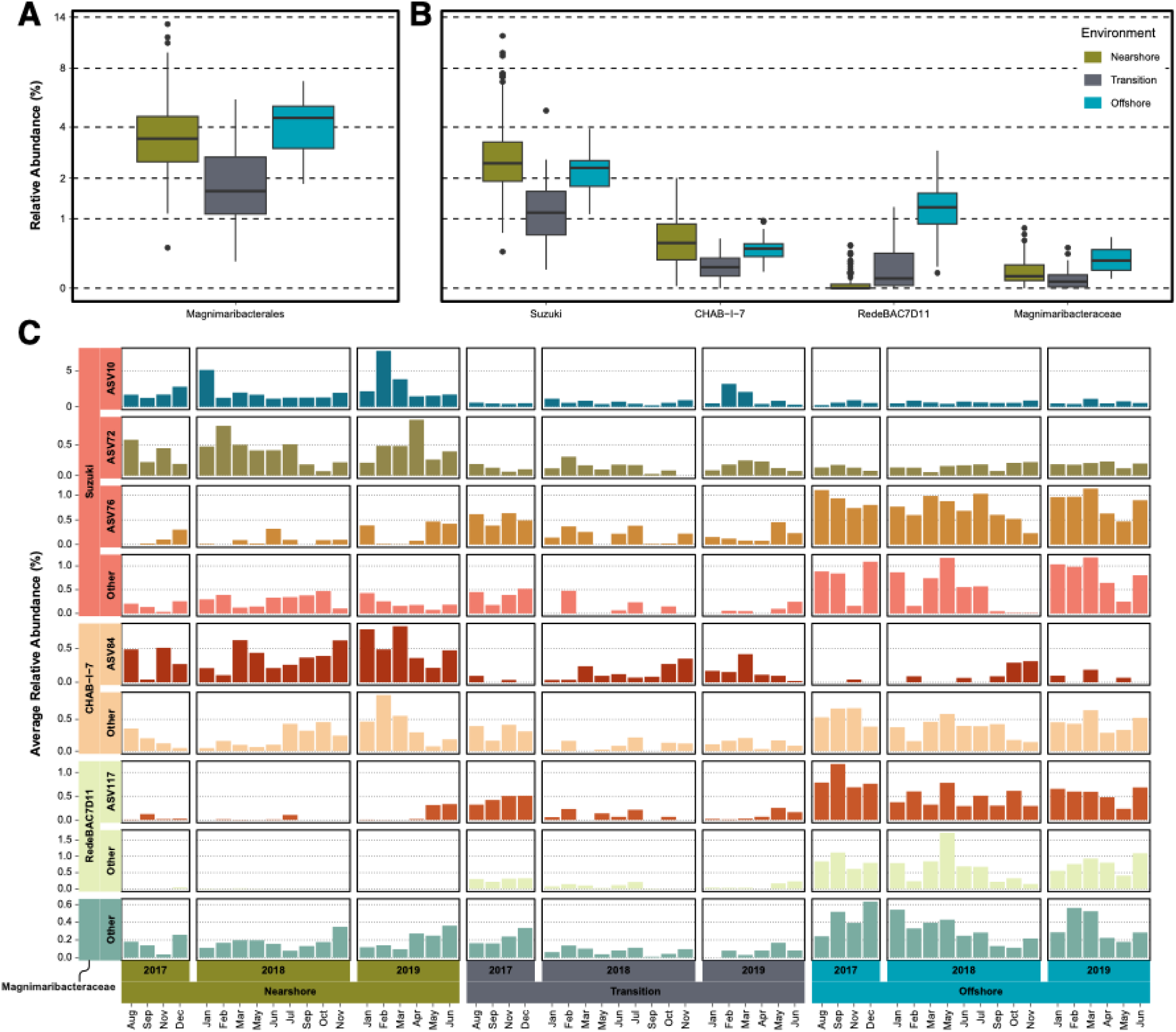
Relative abundance of (A) the order *Magnimaribacterales*, (B) families within the *Magnimaribacterales*, and (C) ASVs within the *Magnimaribacterales* across KByT, based on 16S rRNA gene amplicons. All ASVs that did not exceed 1% relative abundance are grouped as “other”. For (A) and (B), the line in the box displays the median value, hinges indicate the 25th and 75th quartiles, and whiskers indicate the lowest and largest value within a range of 1.5x the interquartile range. Dots indicate outliers.

Of 168 ASVs identified as part of the *Magnimaribacterales* order (Figure S1), only 38 were found in more than one sample and had more than 10 reads associated with them (Table S5). Of the 38, five were frequently responsible for a majority of the *Magnimaribacterales* relative abundance, especially in the nearshore environment (Figure S4). These include three Suzuki, one CHAB-I-7, and one RedeBAC7D11 family ASVs (Table S5).

The five abundant ASVs were not uniformly distributed across the KByT study area (Figure 2C, Figure S4, Table S5). Over the two years of this amplicon dataset, ASV10 and ASV72 from Suzuki and ASV84 from CHAB-I-7 were more abundant in the nearshore environment compared to the offshore, while ASV76 from Suzuki and ASV117 from RedeBAC7D11 were more abundant in the offshore environment (Figure 2C, Figure S4, Table S5). One particular ASV from the Suzuki family (ASV10) dominated the nearshore *Magnimaribacterales* community, peaking at over 8% average relative abundance among nearshore samples collected in February 2019 (Figure 2C). ASV10 was largely responsible for the temporal variability in relative abundance of *Magnimaribacterales* among nearshore samples (Figure S4).

### *Magnimaribacterales* read recruitment across KByT and the surrounding open-ocean is dominated by genomes of the family Suzuki

Nine of 12 initial *Magnimaribacterales* KByT MAGs passed quality control (>50% completion and <7.5% contamination; Table S6) and were included in a phylogenomic analysis with a previously assembled database of global *Magnimaribacterales* genomes (3). All nine MAGs branched within one of the previously delineated families and genera of this order (Figure 3, Table S6).

**Figure 3.**
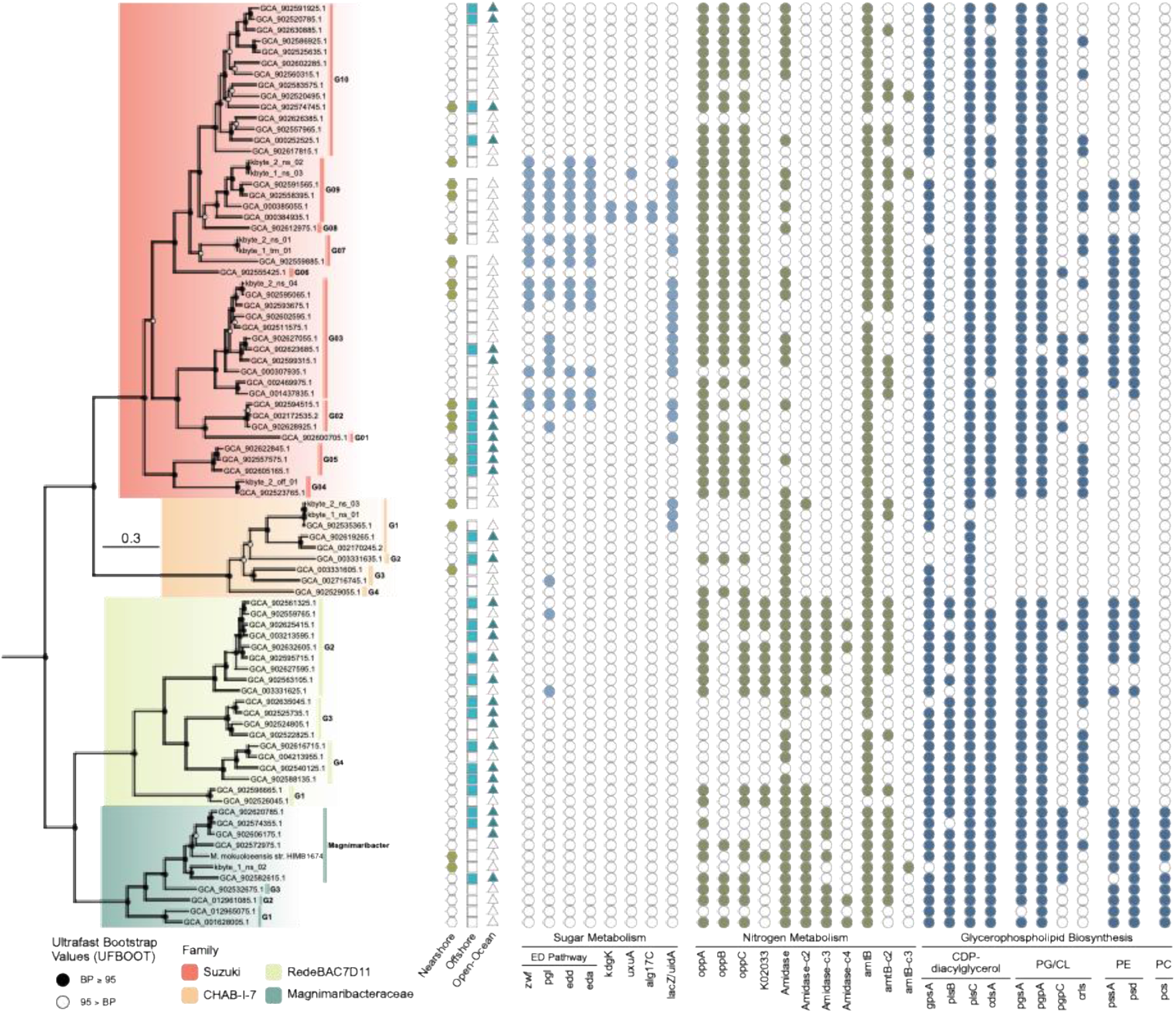
Phylogenomics of the order *Magnimaribacterales* based on a concatenated alignment of 120 single copy core genes. Solid circles indicate bootstrap values greater than 95%, while open circles indicate values over 80%, from 1000 replicates. Four genomes from the gammaproteobacterial orders *Burkholderiales* and *Pseudomonadales* (GCF_000305785.2, GCF_003574215.1, GCF_003752585.1, GCF_006980785.1) were used as an outgroup. Within each family, genera are indicated with the prefix “G”. Filled symbols in the Nearshore, Offshore, and Open-Ocean columns indicate the presence of a genome within each environment based on read recruitment. Genomes without shapes were not recruited since they were too closely related to another genome. For metabolic genes, filled circles indicate the presence of a gene within each genome. Genes prefixed by ”c-” indicate the copy number of those genes for the respective genome. ED - Entner-Doudoroff; PG/CL – phosphatidylglycerol/cardiolipin; PE - phosphatidylethanolamine; PC - phosphatidylcholine.

Consistent with 16S rRNA gene amplicon sequence data, metagenome read recruitment to *Magnimaribacterales* genomes revealed that the vast majority of reads recruited to members of the Suzuki family (Figure 4A, Table S7). Relative to the *Magnimaribacterales* community, families RedeBAC7D11 and *Magnimaribacteraceae* increased in abundance in the offshore and open-ocean environments, with RedeBAC7D11 almost completely absent from the nearshore environment. In contrast, CHAB-I-7 genomes increased in abundance relative to the total *Magnimaribacterales* community in the nearshore environment (Figure 4A, Figure S5)

**Figure 4.**
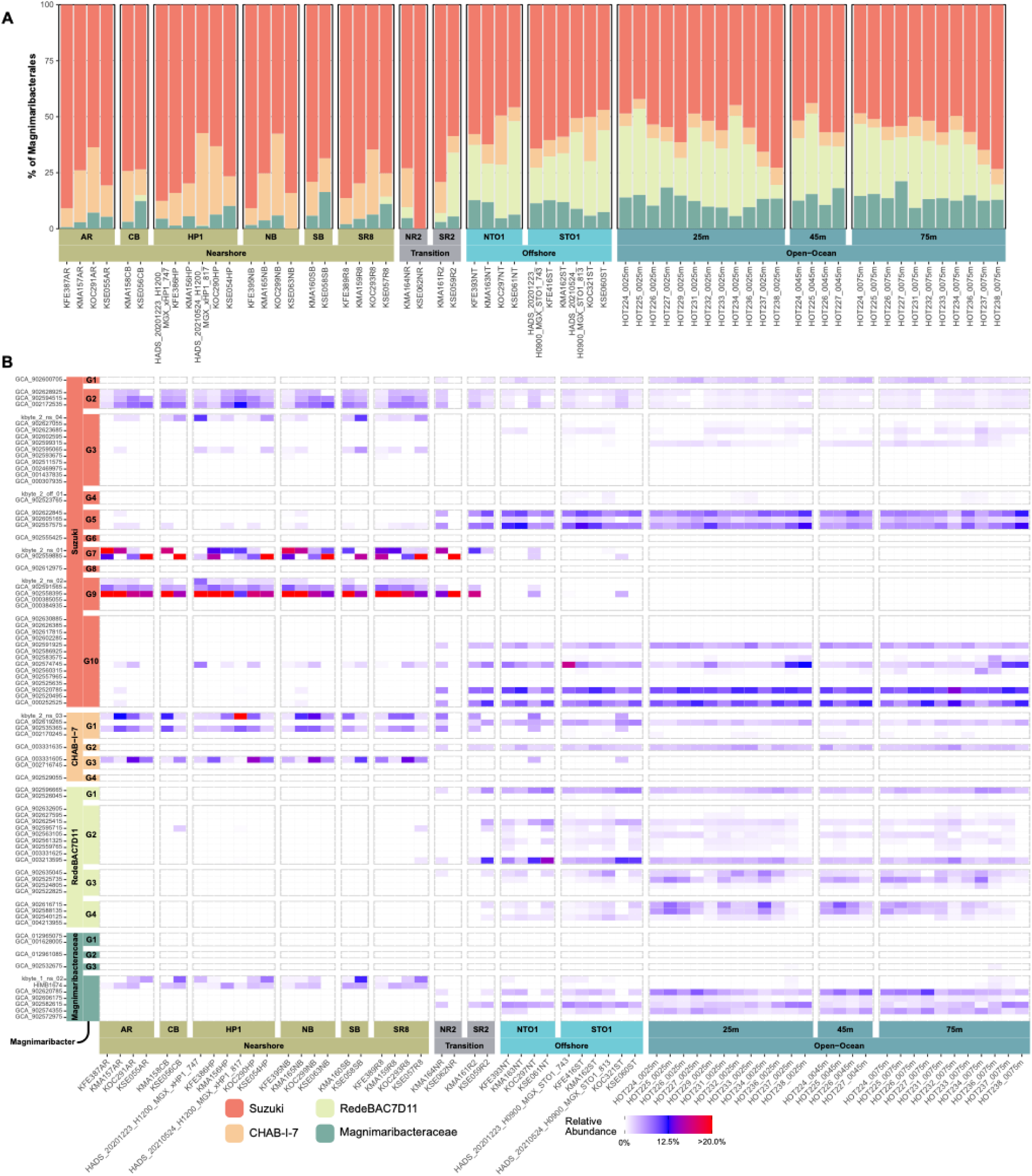
(A) Proportion of the four families to the total *Magnimaribacterales* read recruitment from metagenomes sequenced from KByT and Station ALOHA, (B) Heatmap indicating the proportion of each representative *Magnimaribacterales* genome to the total *Magnimaribacterales* read recruitment within each metagenome.

A small subset of *Magnimaribacterales* genera were responsible for the majority of recruited reads. These genera largely originated from the Suzuki family, where five (G02, G05, G07, G09, and G10) were responsible for >75% of reads recruited to Suzuki and over half of the reads recruited to the *Magnimaribacterales* as a whole (Figure 4, Figure S5). These five Suzuki genera, two (G07 and G09) dominated the recruitment of metagenomes from the nearshore environment, recruiting an average of roughly half of all *Magnimaribacterales* reads (Figure S5). Meanwhile, two other genera (G05 and G10) dominated the recruitment of offshore KByT and open-ocean Station ALOHA metagenomes that together recruited an average of over 40% of total reads (Figure 4, Figure S5). While the genomes within Suzuki genus G02 were most abundant in the nearshore, they did not significantly decline in relative abundance in the offshore to open-ocean environments (Figure S5, Table S8).

Similar to the distribution of 16S rRNA gene amplicons, genomes within the RedeBAC7D11 family were almost entirely limited to offshore and open-ocean environments (Figure 4, Figure S5). However, one genus (G2) was most abundant in the offshore environment immediately outside of Kāneʻohe Bay and declined at open-ocean Station ALOHA, while two others (G3 and G4) were most abundant in the open-ocean environment ( Figure S5, Table S8). Read recruitment to the CHAB-I-7 family revealed genera with distinct nearshore or offshore and open-ocean patterns of distribution. Within the *Magnimaribacteraceae*, members of the genus *Magnimaribacter* were the only genomes consistently detected across all environments, peaking in the open-ocean at Station ALOHA (Figure 4, Figure S5, Table S8).

While the closely related genomes within individual genus-level genome clusters usually followed a similar pattern of distribution, exceptions were evident. At the genus level, CHAB-I-7 genus G1, Suzuki genus G03, and the *Magnimaribacter* genus were broadly distributed across the nearshore to open-ocean environments (Figure 4, Figure S5). However, read recruitment to individual genomes within each of these genera revealed some genomes that were limited to the nearshore environment with closely related neighbour genomes limited to the offshore and open-ocean environments (Figure 4). Read recruitment also revealed temporal differences between the closely related genomes kbyte_2_ns_01 and GCA_902559885 from Suzuki genus G07 (Figure 4B) with the former genome peaking early in the year, while the latter often peaking in September.

Finally, read recruitment was used to categorize each of the 81 genomes as present or absent from each of the nearshore, offshore, and open-ocean environments (Figure 3). Of 42 genomes categorized as “present” in at least one of the environments of this system, 34 were limited in distribution to either the nearshore or offshore and open-ocean environments, five were present across all environment types, and three were present only in the open-ocean environment (Figure 3, Table S3). The five genomes present in all three environments originated from the Suzuki family and primarily from a single genus, Suzuki G02.

### The Entner-Doudoroff pathway is associated with abundant nearshore *Magnimaribacterales*

In general, all families of the *Magnimaribacterales* order contained a complete or near complete Embeden-Meyeroff-Parnas (EMP) pathway for glycolysis except for RedeBAC7D11, which consistently lacked a putative pyruvate kinase gene as has been previously described (Table S9; 3). In contrast to the wide distribution of the EMP pathway among *Magnimaribacterales*, we observed that the full complement of genes for the Entner-Doudoroff (ED) pathway for glycolysis (*zwf*, *pgl*, *edd*, *eda*) was restricted to genomes of a limited number of nearshore-associated lineages within the Suzuki family (G02, G03, G07, G09) (Figure 3, Figure 5).

**Figure 5.**
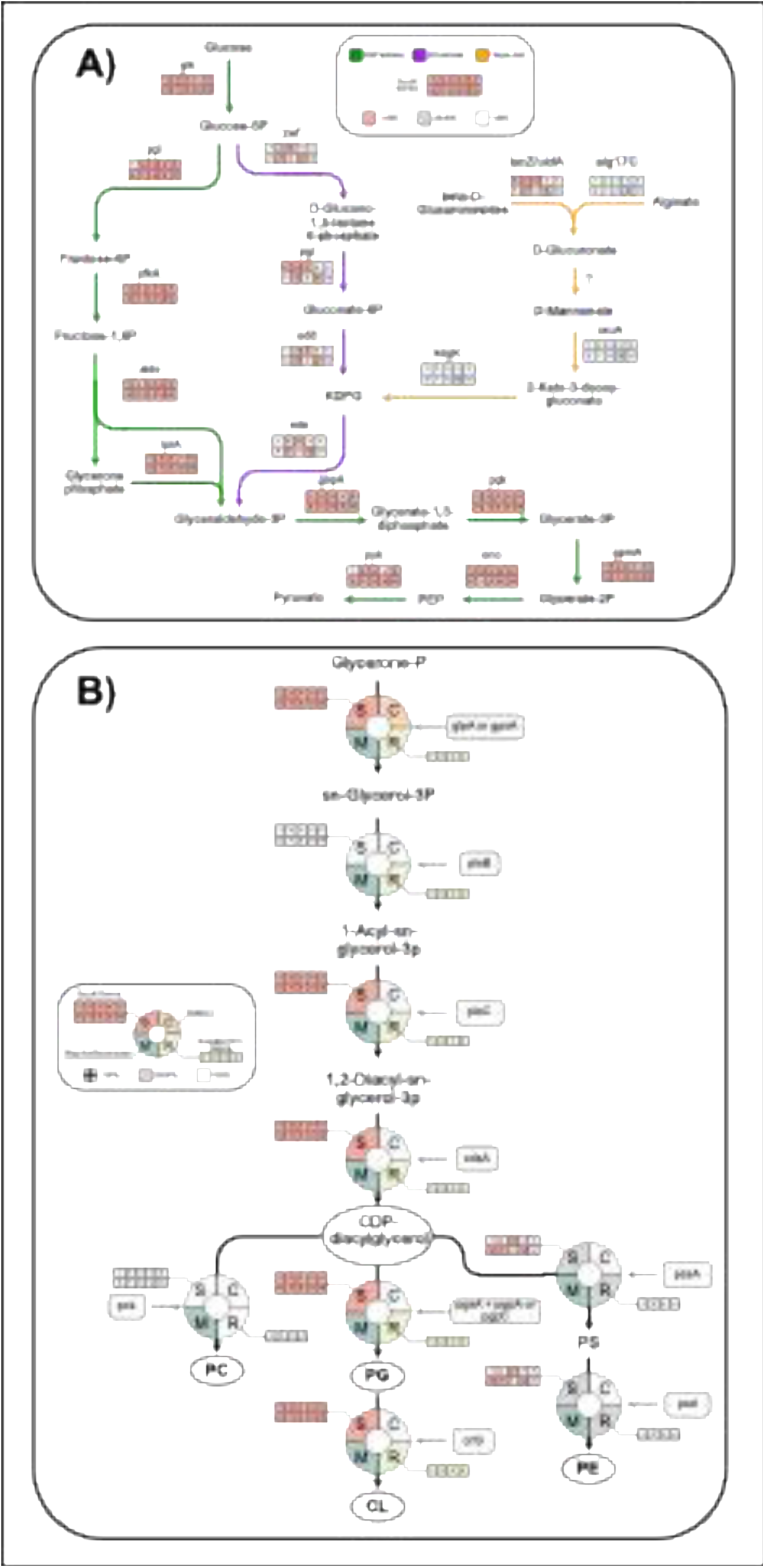
Presence of genes within (A) the Entner-Doudoroff (ED) pathway in genomes of the Suzuki family, and (B) glycerophospholipid biosynthesis pathways in genera of the Suzuki and RedeBAC7D11 families, as well as the families CHAB-I-7 and the *Magnimaribacteraceae*. Genes associated with the ED pathway are based in part on (62), while genes for the synthesis of glycerophospholipids are based on (66). Gene names are indicated in Table S9.

Because the presence of the ED pathway is frequently associated with the metabolism of glucose and various sugar acids, *Magnimaribacterales* genomes were manually inspected for genes related to these functions. Three genomes from Suzuki genus G09 encoded a glucuronate isomerases (*uxaC*), with two of these genomes also encoding a putative 2-dehydro-3-deoxygluconokinase (*kdgk*). These genes function in a pathway that converts the sugar acid D-glucuronate into 2-keto-3-deoxy-6-phosphogluconate (KDPG) (62), which can be subsequently metabolized by the ED pathway (Figure 5). Among the genomes encoding the ED pathway, two potential sources of D-glucuronate were identified: alginate and beta-D-glucuronosides. A small subset of genomes encode putative oligo-alginate lyases (*alg17C*) that are capable of converting alginate into sugar acids (63). Additionally, putative beta-galactosidase/beta-glucuronidases (*lacZ/uidA*) capable of converting beta-D-glucuronosides to D-glucuronate were also present in ED-associated Suzuki genomes (Figure 3). However, three CHAB-I-7 genomes were also found to encode putative copies of this gene. As the CHAB-I-7 family entirely lacked the pathway ED genes, the putative role of this gene as a funnel into the ED pathway is uncertain.

### Differences associated with nitrogen metabolism distinguish closely related phylotypes within the RedeBAC7D11 family

Since genera of the family RedeBAC7D11 differ in their distributions between offshore and open-ocean environments (Figure 4), we next sought to compare the genomes of the four RedeBAC7D11 genera to identify functions associated with these distinct spatial patterns. Enrichment analysis revealed that two oligopeptide transport system genes, *oppA* and *oppC*, distinguished the two offshore RedeBAC7D11 genera (G1 and G2), from the two that peak in abundance in the open-ocean (G3 and G4; Figure 3). An expanded search for the other three opp related genes (*oppB*, *oppD*, and *oppF*; 64, 65) found no homologs in RedeBAC7D11 genomes, though *oppB* along with *oppA* and *oppC* were present in other *Magnimaribacterales* family genomes.

Inspection of the genomic region surrounding the putative opp system genes *oppA* and *oppC* within the RedeBAC7D11 genera G1 and G2 revealed these genes to flank a putative peptide/nickel transport system permease gene (K02033). As RedeBAC7D11 genera G1 and G2 appeared to lack a *oppB*, this gene may support the functions carried out by *oppB*.

To assess whether the absence of the opp system in RedeBAC7D11 G3 and G4 was not simply due to missing contigs, a comparative analysis was performed focusing on the genomic region containing the opp system in RedeBAC7D11 G1 and G2. A putative peptidoglycan lytic transglycosylase D (*mltD*) gene, located near the *oppA* and *oppC* genes, was identified as a consistent feature across all RedeBAC7D11 genomes, each encoding a single copy. This gene was subsequently used to anchor the region of interest. The resulting comparison revealed a homologous genomic segment across all four RedeBAC7D11 genera, displaying a high degree of concordance in both functional annotations and gene synteny (Figure S6).

The transport of oligopeptides suggested by the presence of the opp system in RedeBAC7D11 genera G1 and G2 was associated with differences in the gene repertoire responsible for ammonia transport. Most of the genomes within RedeBAC7D11 genus G2 contained two copies of the ammonia transporter *amtB*, while the RedeBAC7D11 genera G3 and G4 contained only a single copy (Figure 3, Table S9). Of two genomes within RedeBAC7D11 genus G1, one contained two *amtB* homologs while the other lacked genome sequence data that corresponds to this genomic region.

In addition to two *amtB* homologs and the opp oligopeptide transport system, the genomes of RedeBAC7D11 genera G1 and G2 were also enriched in genes encoding for putative amidases (Table S9). While amidases are present across almost all RedeBAC7D11 genomes, the genera G1 and G2 encoded up to two and four homologs respectively, while RedeBAC7D11 genera G3 and G4 generally contain a single copy (Figure 3).

### Pathways for glycerophospholipid biosynthesis distinguish the four *Magnimaribacterales* families

The Suzuki, RedeBAC7D11, and *Magnimaribacteraceae* families broadly encode pathways necessary to biosynthesize the glycerophospholipids phosphatidylglycerol (PG) and cardiolipin (CL) (Figure 5, Table S9). In contrast, the CHAB-I-7 family appears to entirely lack the ability to synthesize glycerophospholipids *de novo*. Additionally, the family lacks many of the genes required to synthesize CDP-diacylglycerol, a key glycerophospholipid precursor (Figure 3).

Genes associated with the biosynthesis of a third glycerophospholipid, phosphatidylethanolamine (PE) (66), were more limited. These genes were present in the *Magnimaribacteraceae* family and a subset of genera within the Suzuki and RedeBAC7D11 families (Figure 3, Figure 5, Table S9). The Suzuki genera capable of synthesizing PE broadly overlap with those that are abundant in the nearshore environment (genera G03, G06, G07, and G09; Figure 3). Within the RedeBAC7D11 family, the pathway for PE biosynthesis was limited to RedeBAC7D11 genus G2, that is more abundant in the offshore compared to the open-ocean environment (Figure 3, Figure S5, Table S8). The gene, *pcs*, supporting the synthesis of a fourth glycerophospholipid, phosphatidylcholine (PC), were only present in genomes of the *Magnimaribacteraceae* family.

## Discussion

Despite three decades of documenting the prevalence of *Magnimaribacterales* bacteria across the global surface ocean (1–3, 5, 67), the nature and extent of ecotypic differentiation within this genetically diverse order-level group remains poorly understood. Building upon our previous work that established a highly curated phylogenomic framework for the *Magnimaribacterales* (3), we leveraged environmental, genomic, and metagenomic data from the geographically constrained Kāneʻohe Bay Time-series (KByT) study system and adjacent Hawaiʻi Ocean Time-series (HOT) study site at Station ALOHA in the North Pacific Subtropical Gyre (NPSG) to uncover how ecotypic differentiation shapes the diversity of this globally abundant order. The results presented here show that *Magnimaribacterales* families, genera, and individual genomes partition across the nearshore to open-ocean waters of the tropical Pacific, with ecotypic differentiation emerging episodically throughout the evolutionary history of this lineage. *Magnimaribacterales* genomes present in this system are largely divided into nearshore and open-ocean specialists, with only a small number of generalist genomes present throughout the entire gradient. Putative functional traits of genomes reflected these spatial differences, as independent lineages occupying the same spatial distribution showed overlap in enriched genes. These findings underscore the importance of ecotypic differentiation in understanding the evolution and diversity of abundant marine bacteria like the *Magnimaribacterales*.

The paraphyletic distribution of nearshore-associated genera across the *Magnimaribacterales* phylogeny, where these genera are interspersed with other differentially distributed genera, may be indicative of a reduced capacity for more closely related genera to coexist within this environment. Indeed, this pattern aligns with recent reports of overdispersion within other abundant marine such as *Pelagibacterales* (11), where lineages more distantly related to one another are more likely to be able to coexist (68). The drivers of the observed overdispersion remain unknown. However, two potential explanations include the competitive exclusion of closely related lineages, or the convergence of function among more distantly related lineages to a similar niche (69). Convergence on function is an unlikely explanation for the phylogenetic overdispersion examined here because, while this study identified several enriched genes, these genes were limited in distribution to only within the Suzuki family.

Instead, the disparate distribution of the nearshore-associated lineages across the *Magnimaribacterales* coupled with the limited overlap in enriched functions across these lineages point towards competitive exclusion as a more likely driver of overdispersion for the broad scale patterns of *Magnimaribacterales* we observe within this system.

The overlap in enriched functions among nearshore-associated Suzuki genera may provide evidence of how environmental drivers shape the community of Suzuki. The association between the nearshore environment and the suite of genes for the synthesis of different glycerophospholipids within Suzuki genomes suggests that these genes may offer some advantage in the nearshore environment and are thus selected for within nearshore-distributed Suzuki genomes. The capacity to vary the lipid composition of the cell has been associated with genome streamlining and the efficient allocation of limited phosphorus and nitrogen available to marine bacterial cells (70–72). The limited diversity of glycerophospholipids that open-ocean-associated genera from the Suzuki family appear able to produce would be indicative of this effect. Conversely, abilities of nearshore-associated Suzuki lineages to biosynthesize PE provide evidence of differentiation in response to an environment where the presence of PE-containing lipids provides a selective advantage. The presence of PE has been shown to play a key role in supporting the function of membrane proteins involved in sugar transport (73). Although this remains speculative, the co-occurrence of the ED pathway, specialized in sugar metabolism, and PE, which may facilitate efficient sugar transport, suggests a linkage between these pathways. These two sets of genes might therefore provide improved fitness in the nearshore environment. Furthermore, the absence of PE has been found in other bacteria to lead to greater sensitivity to osmotic and oxidative stress indicative of the diverse set of functions it influences (74).

Assuming that *Magnimaribacterales* cells remain viable despite an inability to synthesize PE, as has been previous found for other bacteria (73), it does raise intriguing questions regarding the ability of marine bacteria to adapt to the absence of crucial glycerophospholipids like PE. However, the testing of hypotheses related to the presence and absence of PE would greatly benefit from cultivated isolates that could validate the putative functions of the enriched genes, and the impact of their gain or loss on the fitness of cells. Complete genomes from cultivated isolates would also potentially help understand the evolutionary origin of the enriched genes, be it ancestral to the *Magnimaribacterales* order or the Suzuki family or alternatively if they have were acquired through some form of gene acquisition.

This study also provides ample evidence for the ecotypic differentiation of genomes at finer scales than the genome clusters we refer to as genera, indicating likely species-level discrimination within the *Magnimaribacterales*. The temporal variation observed within genera such as Suzuki genus G07, along with spatial differences within the genera *Magnimaribacter* and Suzuki G03, suggest that even subtle genomic distinctions have led to adaptations to specific spatial or temporal niches, differentiating them from closely related genomes. This partitioning of distributions among closely related genomes mirrors patterns seen in other abundant marine bacteria including *Prochlorococcus* (75) and the *Pelagibacterales* (11). A particularly striking example of spatial differentiation was observed in genus G2 of the RedeBAC7D11 family, where several closely related genomes each displayed distinctive patterns of distribution that were often dissimilar to that of their closest genome relative. Similar phenomena have been observed in other marine bacteria, such as comparative analyses of closely related genomes of *Prochlorococcus* and *Pelagibacterales* (13, 54, 76). For example, comparisons of *Prochlorococcus* and *Pelagibacterales* populations between the NPSG and the North Atlantic have revealed biogeographical differences among closely related populations that appear to be associated with only a small set of genes (76). These genes appear heavily driven by environmental variability, including phosphate availability. Other analyses comparing closely related *Prochlorococcus* genomes have found differentiating genes to be located within genomic islands (54). Additionally, recent investigations into *Pelagibacterales* have revealed similar ideas with only a small set of genes appearing to drive distribution differences (13). However, in this case the genes driving these patterns were instead associated within the genomic backbone. Our findings indicate that, similar to other abundant marine bacterial clades, even minimal genetic differentiation can drive the formation of distinct ecotypes within even closely related genome clusters of the order *Magnimaribacterales*.

Coupled with recent studies that delineated ecotypes of *Pelagibacterales* that partition across the nearshore and offshore waters of KByT (11–13), our findings suggest that the evolutionary trajectories of oligotrophic marine bacteria are shaped, at least in part, by the nearshore-offshore environmental gradient and that environmental selection likely plays a key role in partitioning, maintaining, and generating diversity. The insights presented here highlight the value of leveraging time-series data spanning natural environmental gradients to investigate ecotypic differentiation and the evolution of marine bacteria. Incorporating both temporal and spatial dimensions of the data allow for a more comprehensive view of the evolution of *Magnimaribacterales* in this system and permits us to elucidate ecotypes using both time and space. However, we acknowledge that analyses examining a longer span of time would help further disentangle transient events and seasonal trends. Coupled with more cultivated isolates and complete genomes from these isolates would support the investigation of finer-scale patterns that may be observed. These efforts would help to further extend the understanding of the evolutionary forces that have driven the diversity of *Magnimaribacterales* observed here, and more generally how these drivers shape the diversity observed in marine microbial communities.

## Acknowledgements

We thank J. Jones, C. Ratum, H. Mochimaru, A. Boettiger, A. Deck, J. Hunckler, C. Spotkaeff, C. Sullivan, E. Monaghan, E. Kiefl, J. Fuessel, and E. Freel for their assistance collecting samples, G. Robertson for assistance with sonde calibration, and K. Selph for flow cytometry measurements. This research was supported by funding from the National Science Foundation to MSR (grants OCE-1538628 and OCE-2149128), a Data Science Graduate Fellowship to OR (grant OAR-2118222 to G. Jacobs), a Denise B. Evans Fellowship to OR, and the Colonel Willys E. Lord, DVM & Sandina L. Lord Scholarship to OR, and the J. Watumull Merit Scholarship to OR. This is SOEST contribution xxx and HIMB contribution xxx.

## Supplementary Table Legends

Table S1. Summary of samples used in this study.

Table S2. Results of quality filtering on metagenomes used in this study.

Table S3. Relative abundance of *Magnimaribacterales* genomes across the metagenomes used in this study, as a proportion of the total *Magnimaribacterales* metagenome read recruitment.

Table S4. Average, maximum and minimum relative abundances of *Magnimaribacterales* and associated families across nearshore, transition, and offshore environments of KByT, based on 16S rRNA gene amplicon analyses. Also included are estimated average cellular abundances for *Magnimaribacterales* families, including the standard deviation for each mean in parentheses.

Table S5. Average relative abundance of *Magnimaribacterales* ASVs across KByT based on 16S rRNA gene amplicon analyses. ASV categories as in Figure 2C.

Table S6. Characteristics of *Magnimaribacterales* MAGs generated in this study.

Table S7. Average relative abundance of families within the *Magnimaribacterales* based on read recruitment to each metagenome used in this study. Relative abundances are as a proportion of total *Magnimaribacterales* metagenome read recruitment.

Table S8. Average relative abundance of *Magnimaribacterales* genera as a proportion of the total *Magnimaribacterales* community across the 10 KByT stations and three different depths at Station ALOHA in the open-ocean (25, 45, and 75 m). Values within the brackets indicate maximum and minimum relative abundances.

Table S9. Table of information regarding enriched and associated genes investigated in this study. Columns ending with average indicate the average number of gene copies found within that genus and columns ending with proportion indicate the proportion of genomes within each *Magnimaribacterales* genus that contained at least one copy of the associated gene.

## Supplementary Figures

**Figure S1.**
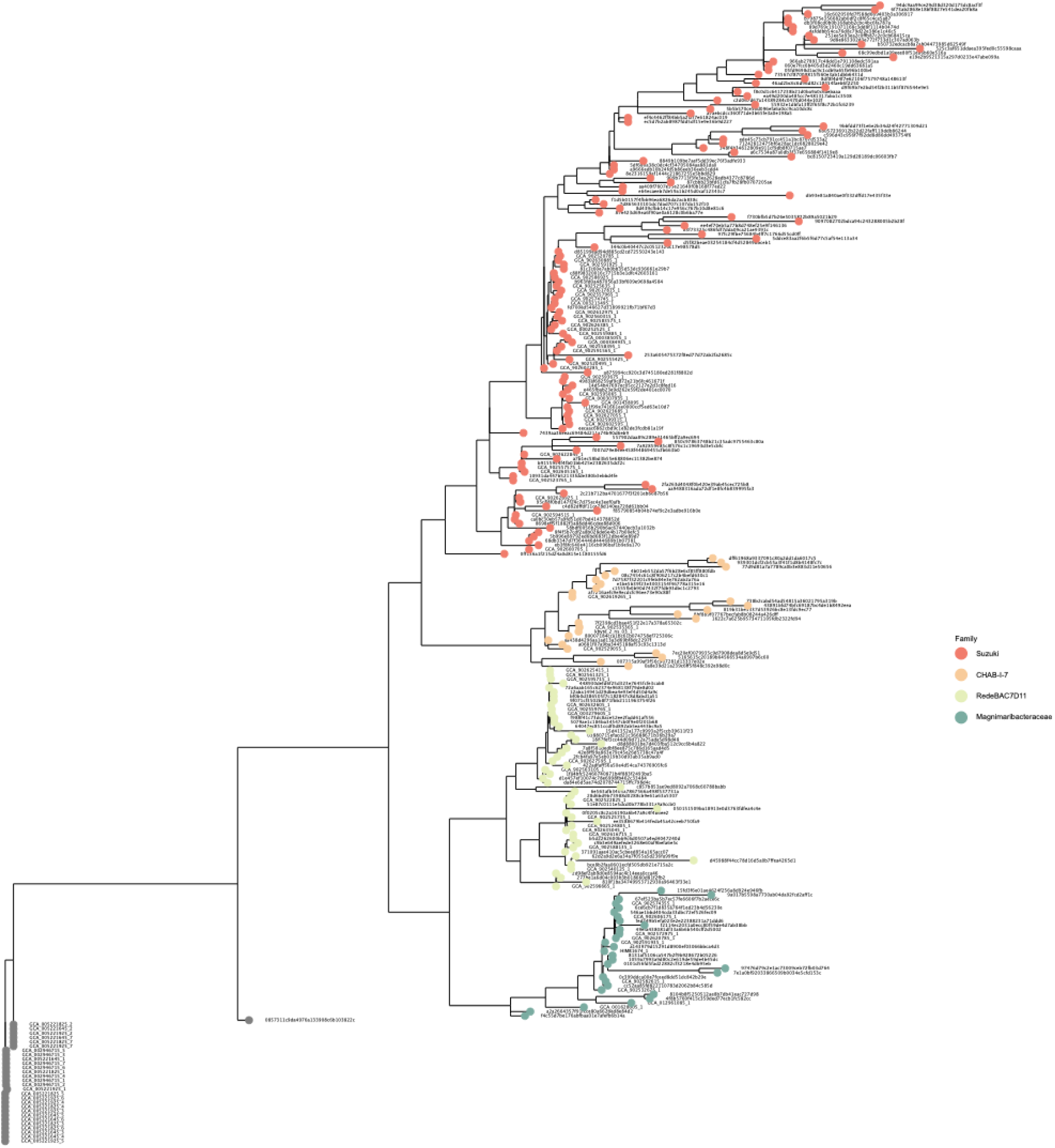
Phylogeny of 16S rRNA gene amplicon ASVs generated from KByT data and 16S rRNA genes from reference *Magnimaribacterales* genomes. The ribosomal RNA genes from four references (GCF_000305785.2, GCF_003574215.1, GCF_003752585.1, GCF_006980785.1) were used as an outgroup.

**Figure S2.**
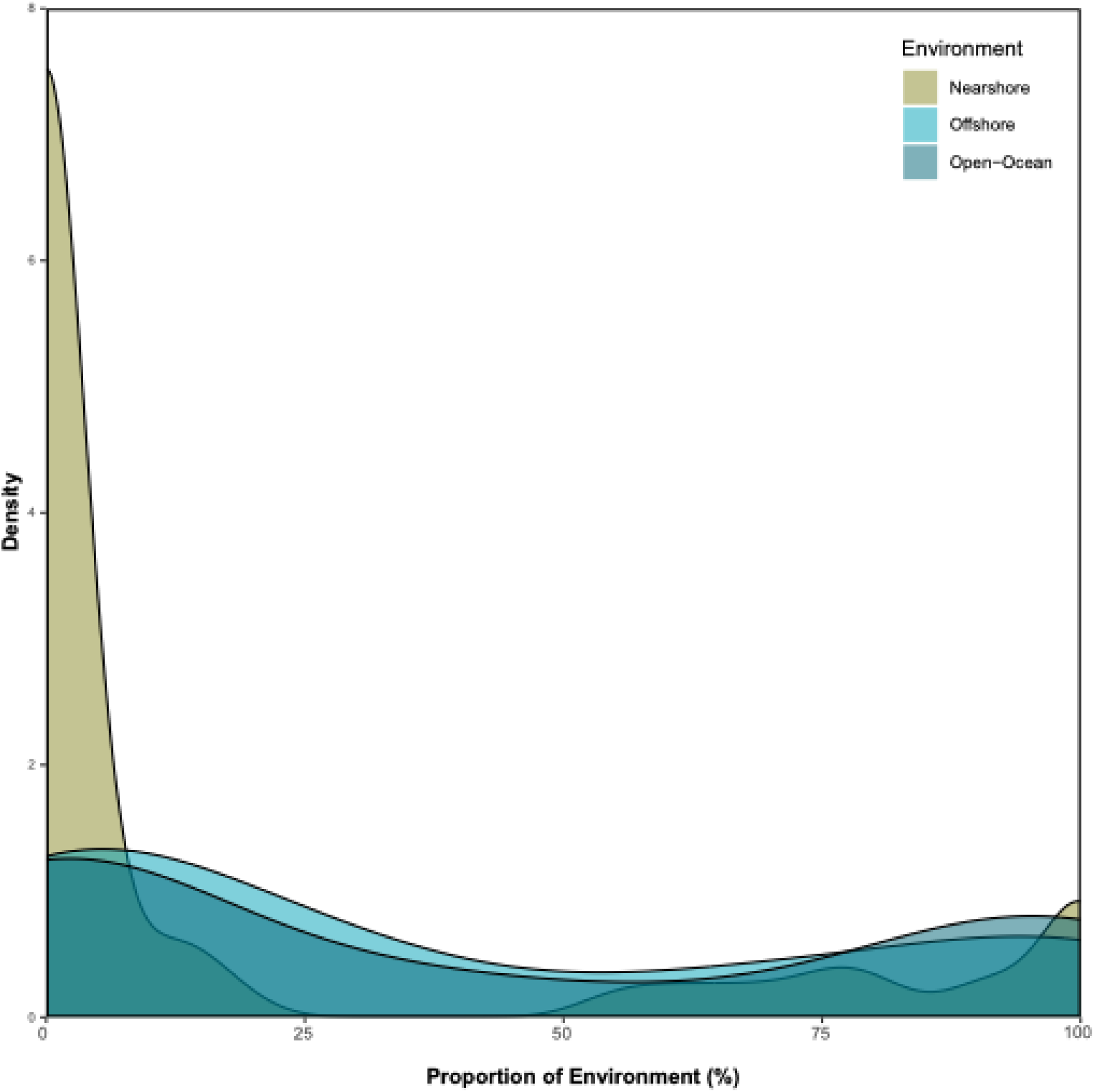
Smoothed histogram showing the distribution of the proportion of metagenomes each genome was detected, separated based on, nearshore, offshore, and open-ocean environments.

**Figure S3.**
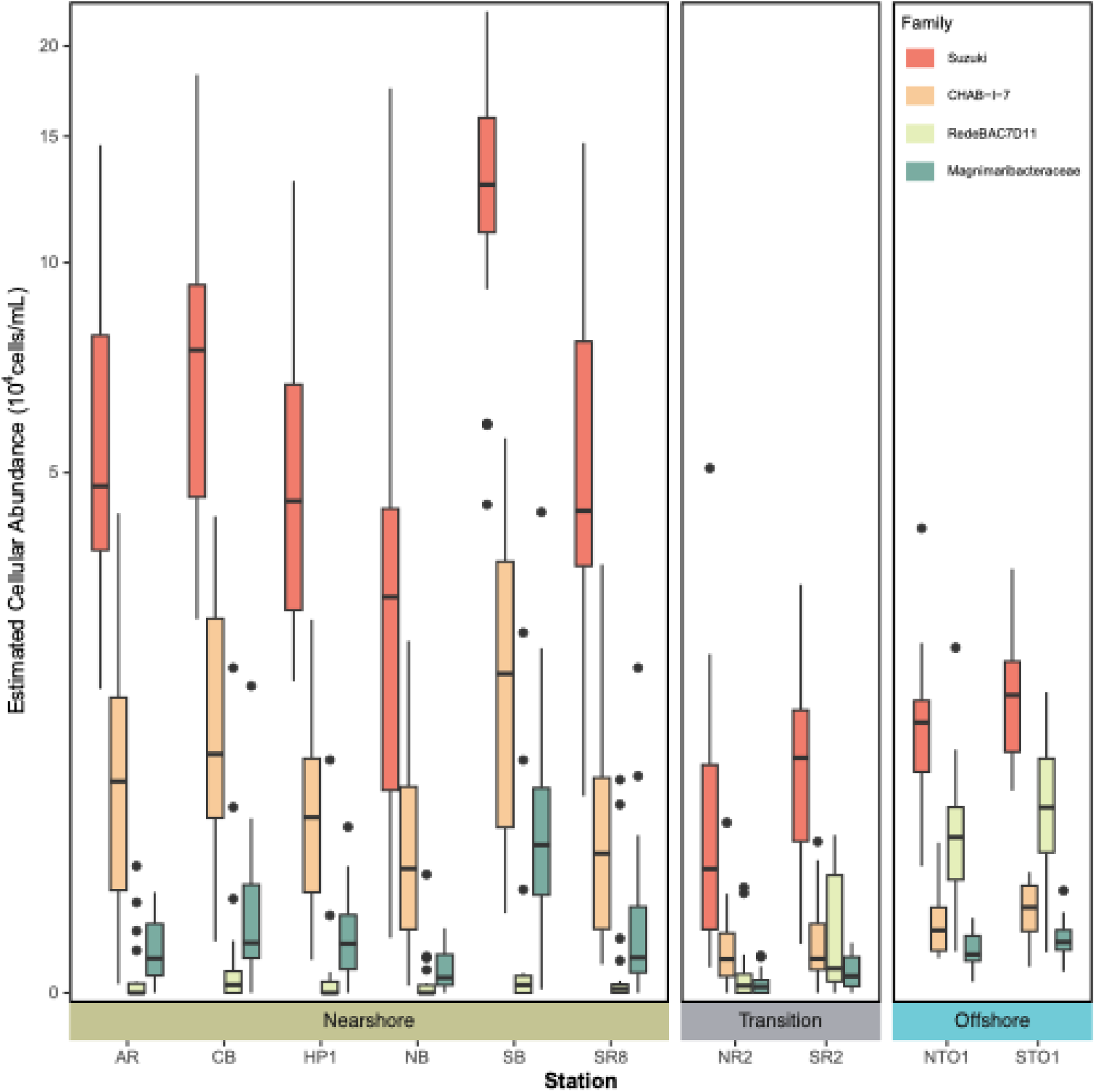
Boxplot showing the estimated cellular abundance of *Magnimaribacterales* families based on the relative abundance of *Magnimaribacterales* to the overall heterotrophic bacteria community and direct cell counts.

**Figure S4.**
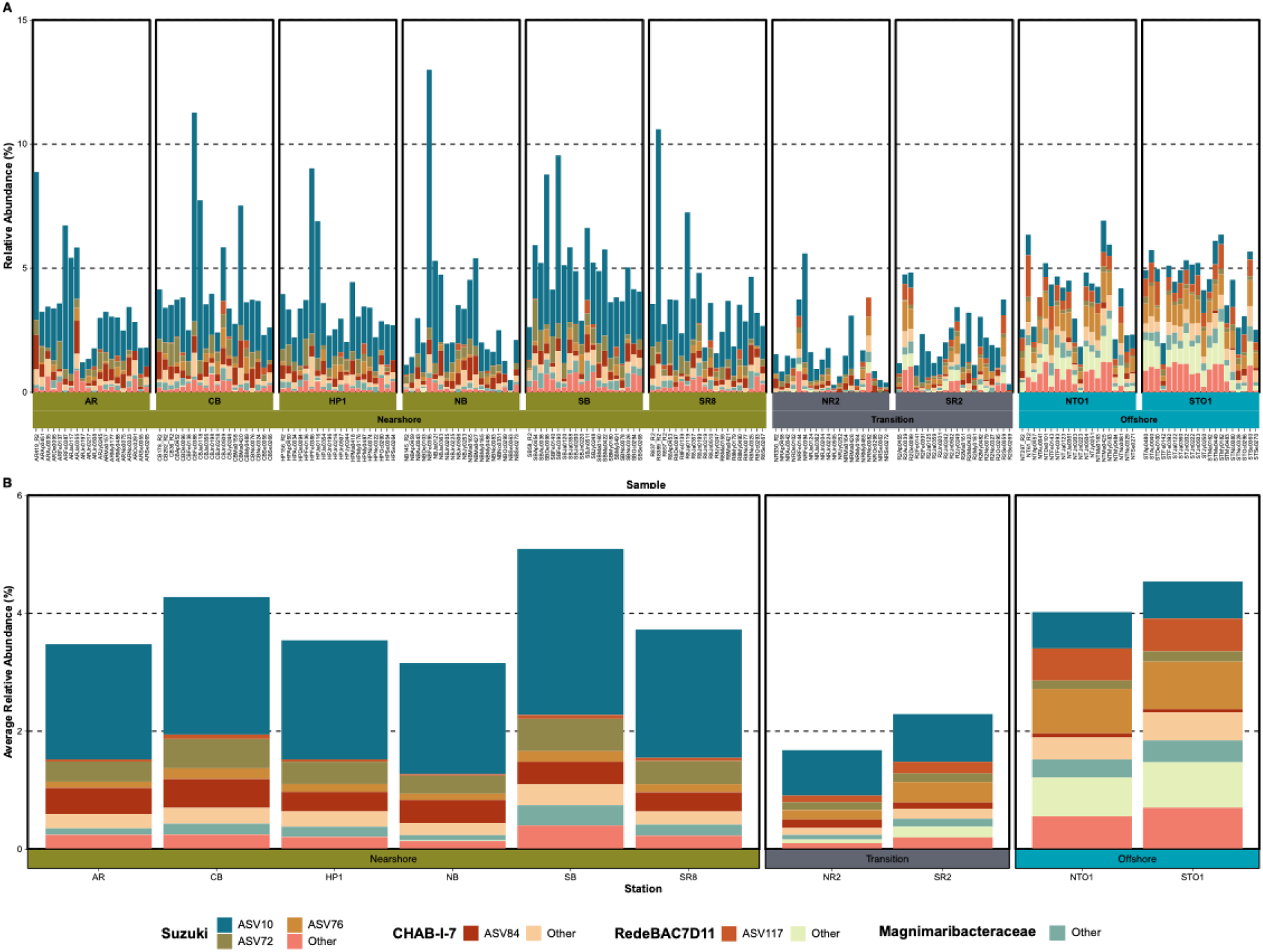
(A) Bar plots of the relative abundance of *Magnimaribacterales* ASVs within each sample, and (B) Average relative abundance of ASVs at each KByT station.

**Figure S5.**
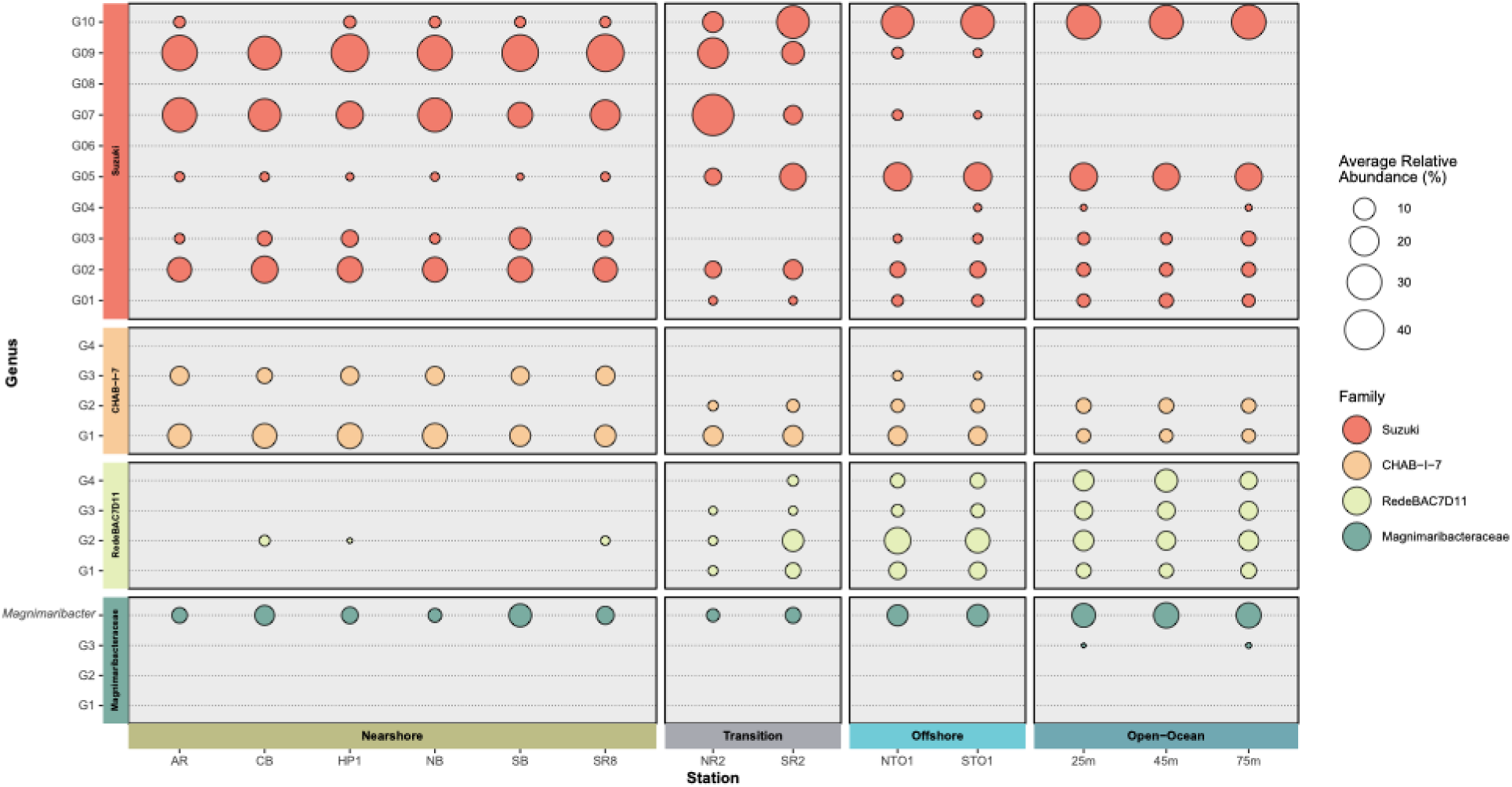
Average relative abundance as a proportion of total *Magnimaribacterales* metagenome read recruitment to genomes of all *Magnimaribacterales* genera across KByT and Station ALOHA in the nearby open-ocean.

**Figure S6.**
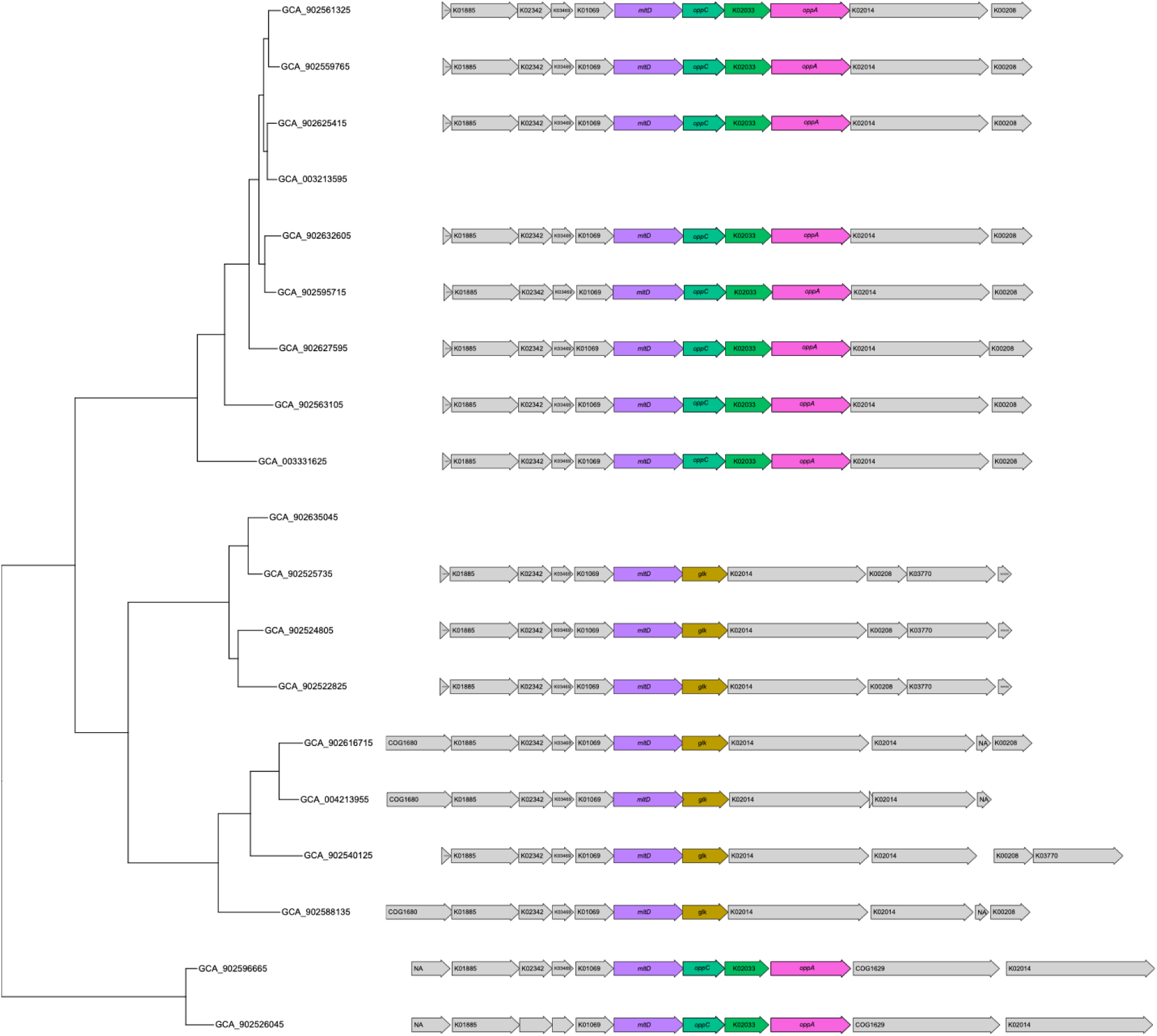
Phylogeny and associated genomic region surrounding the *mltD* gene in genomes from the RedeBAC7D11 family. Colored arrows indicate genes of interest, including putative: peptidoglycan lytic transglycosylase D (*mltD*), glucokinase (*glk*), and the two oligopeptide transport system genes *oppA* and *oppC*, and a peptide/nickel transport system permease related gene (K02033).

